# Systematic analyses of the resistance potential of drugs targeting SARS-CoV-2 main protease

**DOI:** 10.1101/2023.03.02.530652

**Authors:** Julia M. Flynn, Qiu Yu J. Huang, Sarah N. Zvornicanin, Gily Schneider-Nachum, Ala M. Shaqra, Nese Kurt Yilmaz, Stephanie A. Moquin, Dustin Dovala, Celia A. Schiffer, Daniel N.A. Bolon

**Affiliations:** Department of Biochemistry and Molecular Biotechnology, University of Massachusetts Chan Medical School, Worcester, MA, USA; Novartis Institute for Biomedical Research, Emeryville, CA, USA

## Abstract

Drugs that target the main protease (M^pro^) of SARS-CoV-2 are effective therapeutics that have entered clinical use. Wide-scale use of these drugs will apply selection pressure for the evolution of resistance mutations. To understand resistance potential in M^pro^, we performed comprehensive surveys of amino acid changes that can cause resistance in a yeast screen to nirmatrelvir (contained in the drug Paxlovid), and ensitrelvir (Xocova) that is currently in phase III trials. The most impactful resistance mutation (E166V) recently reported in multiple viral passaging studies with nirmatrelvir showed the strongest drug resistance score for nirmatrelvir, while P168R had the strongest resistance score for ensitrelvir. Using a systematic approach to assess potential drug resistance, we identified 142 resistance mutations for nirmatrelvir and 177 for ensitrelvir. Among these mutations, 99 caused apparent resistance to both inhibitors, suggesting a strong likelihood for the evolution of cross-resistance. Many mutations that exhibited inhibitor-specific resistance were consistent with distinct ways that each inhibitor protrudes beyond the substrate envelope. In addition, mutations with strong drug resistance scores tended to have reduced function. Our results indicate that strong pressure from nirmatrelvir or ensitrelvir will select for multiple distinct resistant lineages that will include both primary resistance mutations that weaken interactions with drug while decreasing enzyme function and secondary mutations that increase enzyme activity. The comprehensive identification of resistance mutations enables the design of inhibitors with reduced potential of developing resistance and aids in the surveillance of drug resistance in circulating viral populations.

## Introduction

The COVID-19 pandemic continues to have broad impacts on human health, economics, and day to day life. Multiple strategies have been implemented to reduce the spread of the virus and disease progression, including mRNA vaccines (Polack et al. 2020), monoclonal antibody therapeutics (Chen et al. 2021), and most recently small molecule direct acting antivirals (Owen et al. 2021; Beigel et al. 2020; Jayk Bernal et al. 2022). However, SARS-CoV-2, the virus that causes COVID-19, is capable of evolving rapidly to adapt to new selection pressures (Wang et al. 2021; Starr et al. 2021). For example, recently evolved omicron variants of SARS-CoV-2 have reduced the protection provided by vaccines as well as the efficacy of monoclonal antibody therapeutics (Liu et al. 2022). Evasion of therapeutic interventions is a hallmark of rapidly evolving diseases including many cancers and viral infections (Housman et al. 2014; Kurt Yilmaz and Schiffer 2021). Evaluating the evolutionary potential of therapeutic approaches can provide a valuable guide for genetic surveillance and treatment choices as well as improving the design of new therapeutics with reduced potential for resistance (Figure 1).

**Figure 1.**
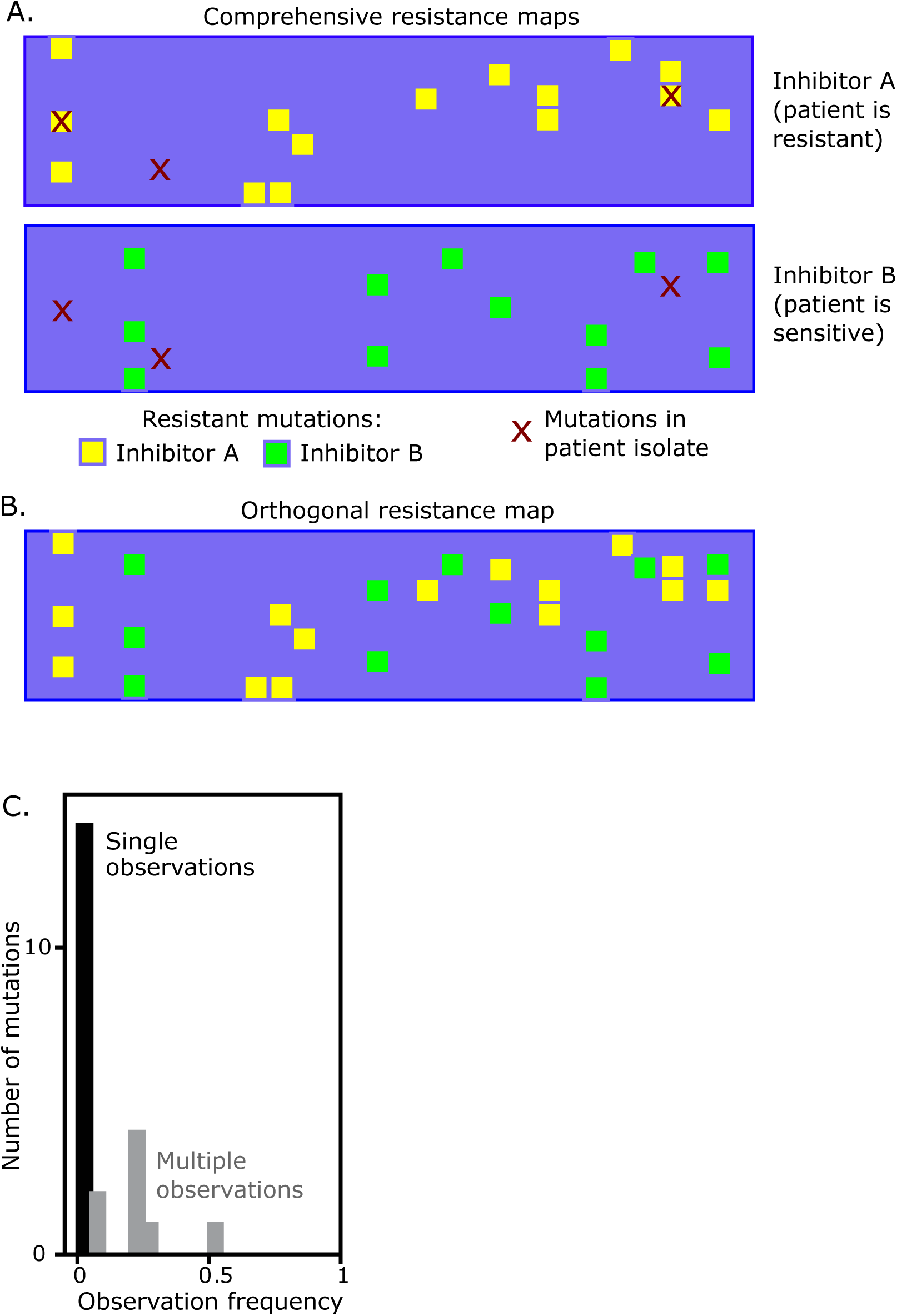
Practical motivations for systematic analyses of drug resistance mutations in SARS-CoV-2 Mpro. (A) Comprehensive maps of drug resistance can enable a sequence-informed choice of inhibitors for effective treatments. (B) Potential to identify inhibitors with orthogonal mutational profiles that could be utilized in combination to reduce the likelihood of resistance evolution. (C) A viral passaging study (Iketani, Mohri et al. 2022) of SARS-CoV-2 reports multiple mutations observed only one time, suggesting that sampling is insufficient to completely reveal all potential resistant mutations. Comprehensive analyses of resistance potential will likely be of use in efforts to survey for evolution of resistance in patient isolates.

Inhibitors targeting the main protease (M^pro^) of SARS-CoV-2 have demonstrated strong clinical effectiveness in treating infected individuals (Owen et al. 2021; Tyndall 2022). The drug PAXLOVID^TM^, a combination of the M^pro^ inhibitor nirmatrelvir and the CYP3A4 inhibitor ritonavir, which boosts the serum half-life of many protease inhibitors, was recently approved for emergency use authorization in treating COVID-19 (Lamb 2022). The wide-scale clinical usage of nirmatrelvir to treat patients infected with SARS-CoV-2 will likely drive selection pressure for resistance; in fact, the evolution of resistance to nirmatrelvir is under active investigation. Several recent reports indicate that multiple mutations in M^pro^ can provide resistance to nirmatrelvir (Iketani et al. 2022b; Jochmans et al. 2023; Hu et al. 2022; Moghadasi et al. 2022; de Oliveira et al. 2022; Zhou et al. 2022; Heilmann et al. 2023). Resistance mutations raised against nirmatrelvir *in vitro* using cell culture exhibit little overlap between different experimental repeats, with many mutations being observed only one time (Figure 1), suggesting that the scale of these traditional approaches is insufficient to provide complete experimental coverage of potential resistance mutations. Additionally, as other M^pro^ inhibitors reach the clinic, it will become important to identify their unique resistance profiles to better guide the development of drugs such that their efficacy is not immediately lost to cross-resistance from prior antivirals (Figure 1).

Analyses of purified M^pro^ harboring drug resistance mutations identified in viral passaging studies indicate that they fall into two categories (Zhou et al. 2022; Jochmans et al. 2023; Iketani et al. 2022a); mutations in the first category weaken binding to drug, while those in the second category increase the proteolytic activity of M^pro^. Similar categories of mutations contribute to drug resistance in many other systems including HIV protease (Ali et al. 2010; Shafer and Schapiro 2008). In HIV protease, a majority of initial mutations arise at the active site, directly affect inhibitor binding, and are the primary cause of resistance to protease inhibitors. These mutations typically also cause reduced enzymatic activity. Subsequently, compensatory mutations arise that rescue the enzyme defects caused by primary mutations; occasionally such mutations appear to also contribute to drug resistance both in HIV and SARS-CoV-2 in cell culture (Iketani et al. 2022b; Zhou et al. 2022; Kozisek et al. 2007; Ragland et al. 2017). In principle, mutations that increase enzymatic activity should provide a small growth advantage in the presence of drugs, and this might be amplified for viral proteases such as M^pro^ which must cleave themselves out of polyproteins to generate additional active protease molecules (Schneider-Nachum et al. 2021). The highest level of drug resistance observed in HIV protease and in SARS-CoV-2 M^pro^ appears to arise from a combination of primary mutations that disrupt drug binding and compensatory mutations that increase enzyme activity.

The potential to evolve resistance to nirmatrelvir provides a strong motivation to comprehensively understand the mutations that can cause resistance. Prior studies in other rapidly evolving diseases have found that identification of drug resistance mutations from clinical samples and/or cell culture passage does not exhaustively identify all resistance mutations (Azam, Latek, and Daley 2003; Kurt Yilmaz, Swanstrom, and Schiffer 2016; Ma et al. 2017; Wagenaar et al. 2014). Here, we describe a comprehensive approach to assess the resistance landscape of M^pro^ from SARS-CoV-2. We provide a systematic map of primary resistance mutations for two chemically distinct inhibitors, nirmatrelvir and ensitrelvir. These two inhibitors exhibit distinct modes of interaction with the M^pro^ active site and we thus hypothesized that they would have distinct drug resistance profiles. We find that these two inhibitors exhibit partially overlapping drug resistance profiles accordant with their chemotypes and interactions in the substrate binding site. We identified both mutations that cause cross-resistance to the two drugs and inhibitor-specific resistance mutations. These comprehensive resistance maps can be used as a guide to evaluate the evolution of drug resistance in circulating SARS-CoV-2 and to aid in the development of drugs with distinct resistance profiles to treat viruses that may evade current drugs.

## Results

### Comparison of the binding modes of M^pro^ inhibitors

We hypothesized that similarities in inhibitor binding interactions may predict similarities in resistance profiles. To investigate this correlation, we compared the interactions of nirmatrelvir and ensitrelvir with M^pro^ to that of other structurally characterized inhibitors. To assess similarities and distinctions in inhibitor binding profiles, we calculated the van der Waals (vdW) interactions between 134 M^pro^-inhibitor crystal structures. This measurement describes the packing of active site residues around individual inhibitors and provides an overall contact pattern for inhibitors. Subsequently, we performed principal component analysis (PCA) to reduce the dimensionality of the data and found that 5 principal components (PCs) accounted for 65% of total variance in the dataset (Figure 2A). M^pro^ inhibitors were grouped into three clusters using k-means clustering on the first five PCs. The three clusters can be differentiated by PC1, which primarily reflects vdW interactions with the inhibitor at the S1 and S2 subsites (where S1 refers to the binding pocket for the substrate amino acid immediately N-terminal to the cut-site, and S2 the binding pocket for the preceding amino acid). Nirmatrelvir and ensitrelvir belong to different clusters (1 and 2 respectively) suggesting that they may also exhibit distinct drug resistance profiles. In future efforts it will be interesting to extensively explore how inhibitor binding classification relates to drug resistance profile.

**Figure 2.**
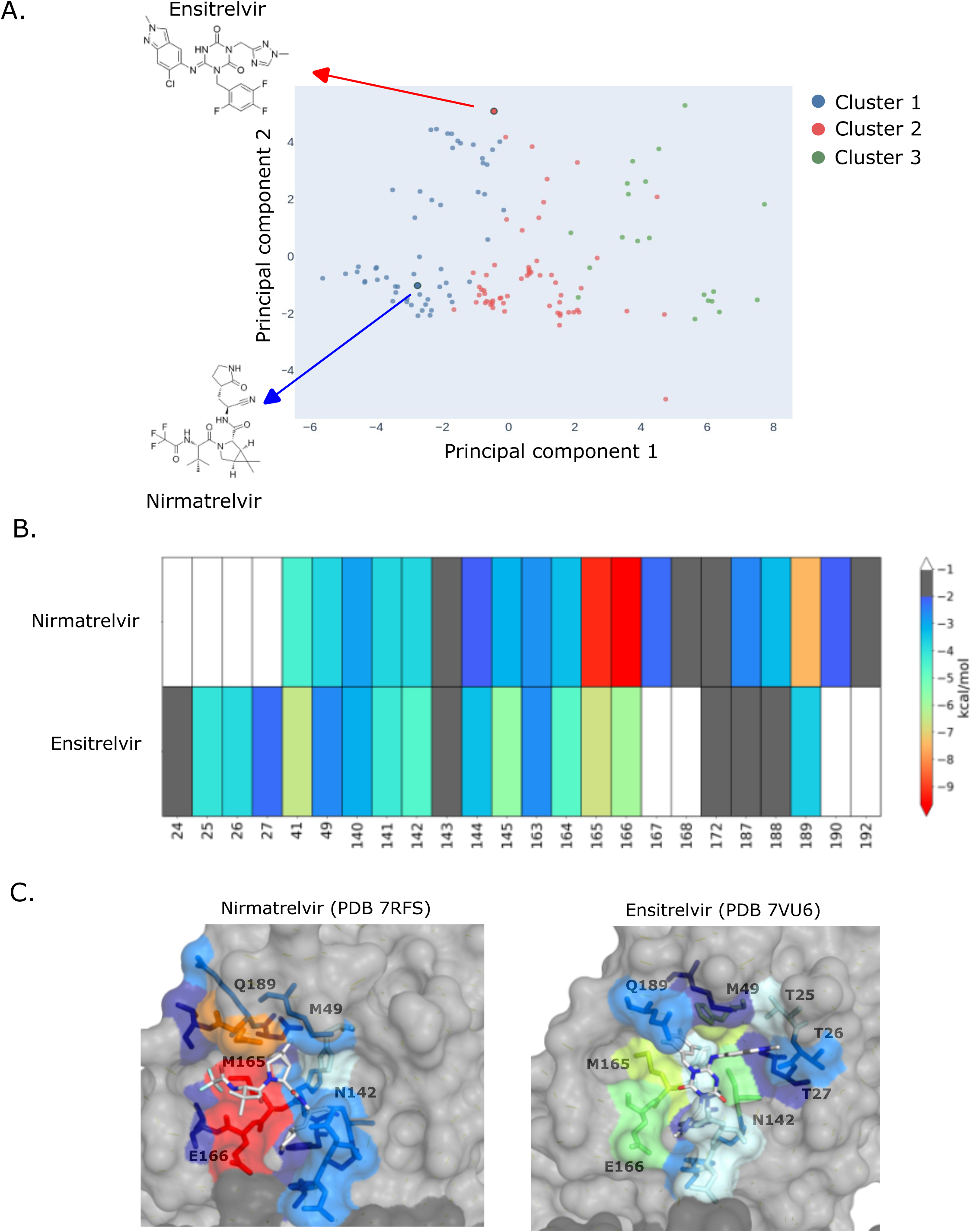
Principal component analysis of vdW interactions show distinct modes of nirmatrelvir and ensitrelvir binding. A) Primary component analysis of potent compounds (IC50 < 1 uM) based on strength of van der Waals interactions. PCA data was grouped on three clusters. Nirmatrelvir and ensitrelvir are in clusters 1 and 2 respectfully. B) Heat map based on strength of van der Waals interactions between inhibitor and Mpro residues. C) Strength of van der Waals interactions between Mpro residues and nirmatrelvir (PDB 7RFS) (Owen, Allerton et al. 2021) or ensitrelvir (PDB 7vu6) (Unoh, Uehara et al. 2022) mapped to structure.

Consistent with their classification in distinct PCA clusters, nirmatrelvir and ensitrelvir contact the M^pro^ active site in distinct modes. Nirmatrelvir spans the S4-S1 portion of the active site and forms an extensive interaction network with the active site involving both vdW interactions with non-prime site residues (i.e.: C-terminal to the cleavage site) as well as hydrogen bonding with H163, E166, and Q189 (Figure 2B and C). In contrast to nirmatrelvir and most other M^pro^ active site inhibitors, ensitrelvir extends into the S1′ pocket and interacts with the threonine cluster (T24-T27). Moreover, its P2 moiety forms a π–π stacking interaction with the catalytic residue H41, displacing it from the active conformation.

### Systematic identification of drug resistance mutations

We previously reported multiple yeast-based approaches to systematically quantify the function of all point mutations in M^pro^ (Flynn et al. 2022). In the current work we utilized the effect of M^pro^ expression on yeast growth as a measure of enzymatic activity. Expression of SARS-CoV-2 M^pro^ reduces the growth rate of yeast cells, but catalytically-inactive variants do not (Flynn et al. 2022; Ou et al. 2022), indicating that changes in growth rate are due to cleavage of endogenous yeast proteins. While this assay relies on cleavage of proteins that may not be directly relevant to SARS-CoV-2, the impacts on yeast growth correlate strongly (R^2^=0.9) with measures from a FRET-based assay that directly measures cleavage of the SARS-CoV-2 Nsp4/5 cut-site (Flynn et al. 2022). In addition, we found that functional scores from yeast growth assays correlated linearly with catalytic rates measured for individual variants (though increased activity variants have not been assessed carefully by alternate approaches). The approach also identified known critical residues (such as catalytic residues), and circulating clinical variants (which must contain active M^pro^) overwhelmingly show wild-type-like functional scores (Flynn et al. 2022). Together, these results suggest that our yeast growth assay provides useful estimates of M^pro^ function for the purpose of assessing SARS-CoV-2 fitness.

To systematically identify potential resistance mutations, we performed deep mutational scanning experiments in the presence of nirmatrelvir and ensitrelvir. Because yeast possess numerous drug efflux pumps and are protected by a cell wall, they often require genetic engineering to make them accessible to drugs (Chinen et al. 2016; Suzuki et al. 2011; Rogers et al. 2001). To increase the druggability of the yeast in our assays, we used a strain called Δ4 with four efflux pumps deleted (Δsnq2 Δpdr5 Δpdr1 Δyap1) and added a low concentration of SDS to increase permeability. Under these conditions, we found that addition of either nirmatrelvir or ensitrelvir restored the yeast growth rate that was retarded due to M^pro^ expression (Supplemental Figure 3). In our mutational scanning experiments, we found that addition of either 10 µM ensitrelvir or 20 µM nirmatrelvir to the culture inhibited approximately 50% of the activity of the wild-type M^pro^ in our library. We targeted this level of inhibition to provide a more sensitive readout of mutations that counteract drug treatment.

We transformed Δ4 strain yeast with our previously described barcoded M^pro^ plasmid library (Flynn et al. 2022) that contains both internal positive controls (wild-type M^pro^) and negative controls (internal stop codons known to destroy function). The resulting yeast libraries were amplified without induction of M^pro^ and subsequently treated with either 20 µM nirmatrelvir, 10 µM ensitrelvir, or DMSO, followed by addition of 2 µM β-estradiol to induce expression of M^pro^. Cells were collected at 0- and 16-hour time points and subjected to Illumina sequencing to estimate the frequency of each variant in each sample (Figure 3A). We calculated a functional score for each variant based on its change in frequency before and after induction of M^pro^ as further described in Materials and Methods (See Supplemental Table 1 for functional scores for each variant). Functional scores were normalized such that a score of 0 represents an inactive mutation that abolishes M^pro^ activity and thus restores yeast growth. Under the conditions of the mutational scans in this work, wild-type M^pro^ had a functional score of 0.67 in the absence of inhibitors, and we were able to reproducibly measure mutations that both increased and decreased enzyme activity. Experiments were performed in triplicate; functional scores between the three replicates were strongly correlated. Figure 3B shows the correlation between the functional scores of all mutations in two replicates measured in the absence of inhibitor (R^2^=0.95) or in the presence of nirmatrelvir (R^2^=0.95) or ensitrelvir (R^2^=0.94).

**Figure 3.**
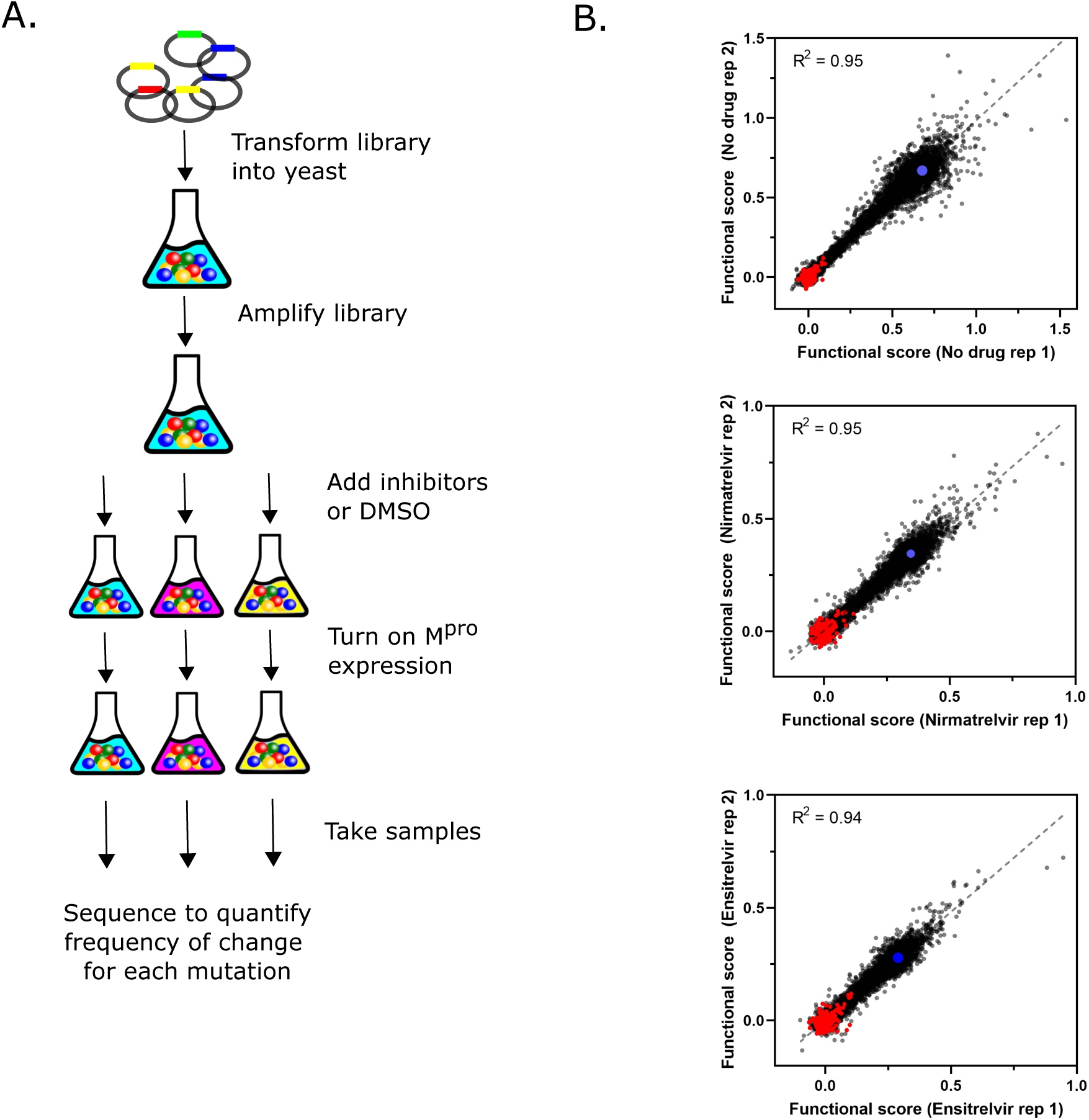
Systematic analyses of mutations that decrease sensitivity to nirmatrelvir and ensitrelvir. A) Illustration of yeast selection screen B) Correlation between replicates of functional scores of all Mpro variants in the absence of drug or presence of nirmatrelvir or ensitrelvir. Stop codons are shown in red and wild-type Mpro in blue.

The distributions of functional scores of all mutations in the presence or absence of inhibitors suggests that both nirmatrelvir and ensitrelvir were effective in reducing the activity of most M^pro^ variants (Figure 4A). Wild-type M^pro^ was inhibited approximately 50% by nirmatrelvir or ensitrelvir, as indicated by the blue dotted lines, and we could clearly distinguish the functional scores of stop codons (shown in red) from wild-type M^pro^ in each experiment (Figure 4A).

**Figure 4.**
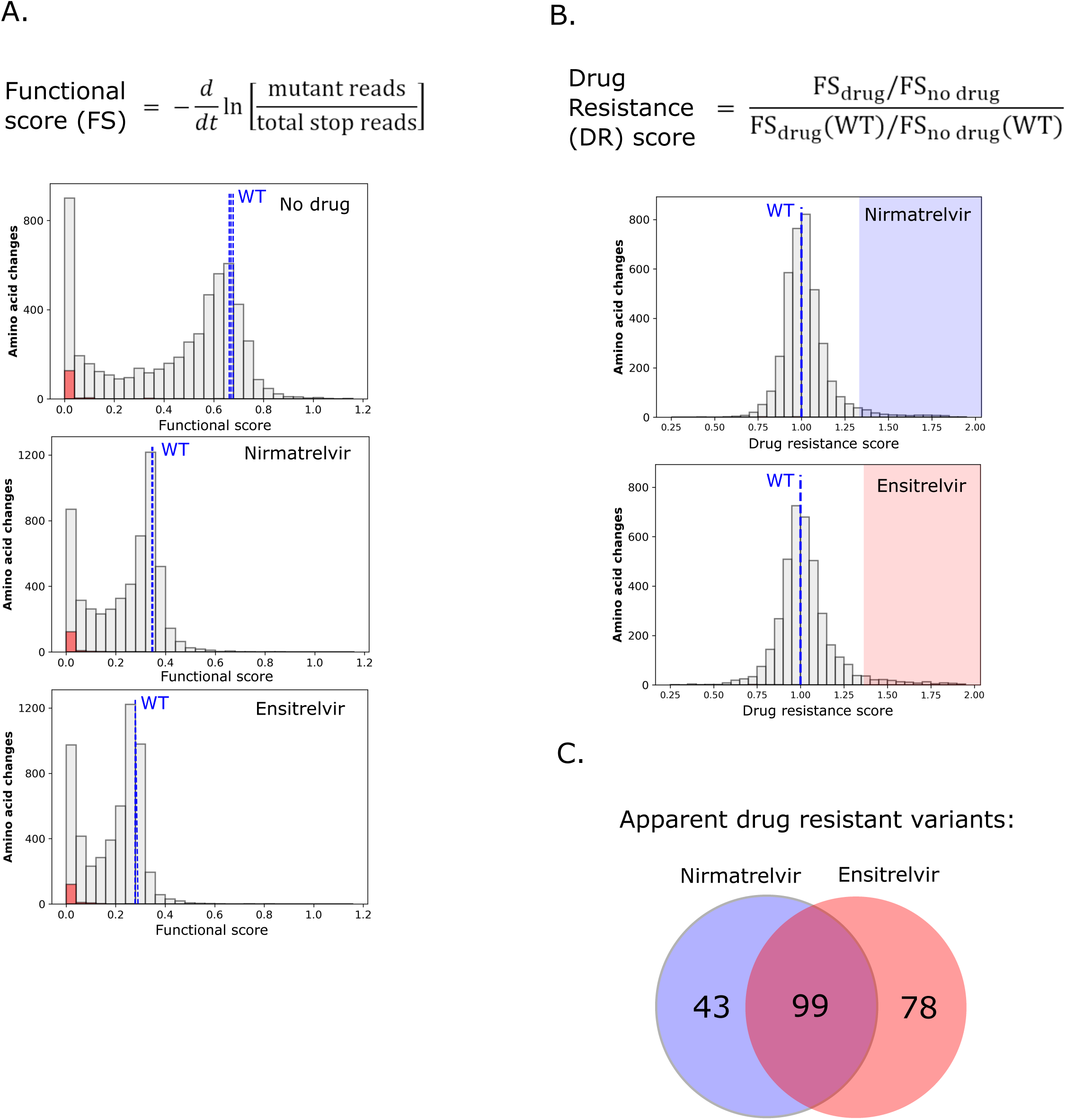
Comprehensive analysis of mutations that show drug resistance. A) Equation to calculate functional scores (top panel). Distribution of all functional scores for all variants (gray), stop codons (red) and wildtype (blue dashed lines) in the absence or drug or presence of nirmatrelvir or ensitrelvir (bottom panel). B) Equation to calculate drug resistant (DR) scores (top). Distribution of all DR scores in the presence of nirmatrelvir or ensitrelvir (bottom panel). Mutants with DR scores above two standard deviations of wildtype are highlighted and are considered apparent drug resistant mutations. C) Venn diagram of nirmatrelvir and ensitrelvir resistant mutations.

Similar to wild-type M^pro^, both inhibitors reduced the functional scores of most mutants by about 2-fold as well, consistent with the conclusion that most mutations did not affect drug resistance. However, a subset of mutations had functional scores that were relatively unaffected by the inhibitors (Supplemental Figure 4A), suggesting these variants may be relevant for drug resistance. To systematically assess the impacts of mutations on inhibition, we calculated a drug resistance score (DR score) (See Supplemental Table 1 for DR scores for all variants). We defined the DR score as the ratio of the functional score of each mutant in the absence of the inhibitor to that in the presence of the inhibitor, compared to the same ratio with wild-type M^pro^ (Figure 4B). By computing the ratio in functional scores with and without drug, the underlying impacts of mutations on enzyme function are canceled out such that the drug resistance score estimates drug sensitivity. This is similar to IC_50_ estimates of drug resistance that quantify changes in sensitivity to drug independently from how a variant impacts function. Under the conditions of our experiments, drug resistance scores have a ceiling of about 2 even if the inhibitors have no measurable impact on function, while wild-type M^pro^ has a DR score of 1. DR scores were only calculated for variants with a functional score at least 1/3 that of wild-type, which approximates the lowest functional score we observed for common, already-circulating M^pro^ variants discussed in our previous work (Flynn et al. 2022). This threshold was applied to limit the potential for elevated noise derived from ratios of small numbers as well as to account for the requirement of M^pro^ activity for viral propagation. Our approach was designed to sensitively identify mutations that cause drug resistance and distinguish the impact these mutations are predicted to have on enzymatic activity.

Figure 4B shows the distribution of DR scores calculated for both nirmatrelvir and ensitrelvir. We considered mutants with DR scores above two standard deviations (SD) of wild-type as apparent drug resistance mutations. We identified 142 mutants with apparent resistance to nirmatrelvir and 177 to ensitrelvir (Figure 4C, Supplemental Figure 4B, Supplemental Table 2). Given the total number of drug resistance measurements made for each inhibitor (3858 variants with function above the 1/3 wild-type threshold), random expectations would lead to 6 variants with apparent resistance to both inhibitors. However, we observed 99 mutations that caused apparent drug resistance to both drugs (p<0.0001, chi^2^), suggesting a largely shared structural basis of drug resistance for both inhibitors.

### Mutations with the greatest likelihood of contributing to drug resistance in nature

Both the likelihood of mutations occurring and their impacts on fitness will determine the likelihood of contributing to drug resistance in nature. While detailed estimates of evolutionary likelihoods will require extensive modeling that are beyond the scope of this work, we have performed analyses of primary factors including the number of nucleotide changes required for an amino acid change and the magnitude of the DR score. Under the recommended treatment regimen of Paxlovid, the concentration of nirmatrelvir is maintained well above that needed to inhibit wild-type M^pro^ (Toussi et al. 2022), providing selection pressure for mutations with large impacts on drug binding. Single nucleotide changes are by far the most common mutation observed in SARS-CoV-2 isolates (Supplemental Figure 4C), consistent with reported mutational biases (Kosuge et al. 2020). In contrast, amino acid changes that cause apparent drug resistance to either nirmatrelvir or ensitrelvir tend to require multiple nucleotide changes, consistent with the greater number of amino acid changes that can be accessed. Multiple nucleotide changes can evolve, especially if they provide a unique fitness advantage. However, in the case of both nirmatrelvir and ensitrelvir, the strongest DR scores of single nucleotide variants are similar to those accessible by multiple nucleotide mutations (Supplemental Table 2). These findings indicate that amino acid changes with the highest DR scores accessible by single nucleotide mutations have the most likelihood of contributing to the natural evolution of drug resistance.

The 16 single nucleotide variants with the strongest DR scores for nirmatrelvir and ensitrelvir are listed in Table 1. Viral passaging with these inhibitors has identified some of the same variants, suggesting that our findings are relevant to viral evolution. For example, H172Y is a mutation that was disclosed by Pfizer to cause strong resistance to nirmatrelvir with a greater than 200-fold increase of K_i_ (Administration 2021). Our data identify M49L and P52S as apparent DR mutations for ensitrelvir and these mutations have also been reported in viral experiments with ensitrelvir (https://www.japic.or.jp/mail_s/pdf/23-11-1-07.pdf). In addition, E166V evolved in multiple lines of SARS-CoV-2 under selection with gradual (2-fold) increases in nirmatrelvir (Iketani et al. 2022b). The evolved line with E166V showed the highest reported drug-resistance with an ability to tolerate roughly 1000-fold higher concentrations of nirmatrelvir than wild-type. Interestingly, many lines evolved with gradual increases in nirmatrelvir concentration did not contain mutations with strong DR scores in our study and instead contained multiple mutations with either small increases in DR score and/or increased functional scores relative to wild-type (Supplementary Table 3). Because the treatment regimen of Paxlovid maintains the concentration of nirmatrelvir 10-fold above the sensitivity of SARS-CoV-2 with wild-type M^pro^, clinical drug pressure may preferentially select for stronger resistance than observed with 2-fold increases in drug concentration in cell culture.

**Table 1.** Single nucleotide mutations with largest nirmatrelvir or ensitrelvir resistance scores.

| Mutation | DR score |  |
| --- | --- | --- |
|  | Nirmatrelvir | Ensitrelvir |
| E166V | 2.0 | 1.8 |
| G2R | 2.0 | 1.9 |
| G2C | 1.9 | 1.7 |
| S139P | 1.8 | 1.0 |
| P168R | 1.8 | 2.2 |
| Q299H | 1.8 | 1.9 |
| G2S | 1.7 | 1.7 |
| N214Y | 1.7 | 1.7 |
| G2A | 1.7 | 1.7 |
| C300R | 1.6 | 1.7 |
| M49L | 1.2 | 2.1 |
| H172Y | 1.5 | 1.9 |
| E166A | 1.6 | 1.8 |
| P52L | 1.0 | 1.8 |
| S139L | 1.6 | 1.8 |
| C44G | 0.9 | 1.8 |

We further examined the validity of our yeast screen by measuring the inhibition of two M^pro^ variants (T25E and M49L) that showed apparent DR specifically for ensitrelvir. Consistent with our DR scores, both variants exhibited increased Ki for ensitrelvir (40-fold for M49L and 300-fold for T25E) compared to wild-type and minimal to no observed impact on Ki for nirmatrelvir (Figure 5). Of note, the T25E amino acid change requires three nucleotide changes, reducing the likelihood of its natural evolution.

**Figure 5.**
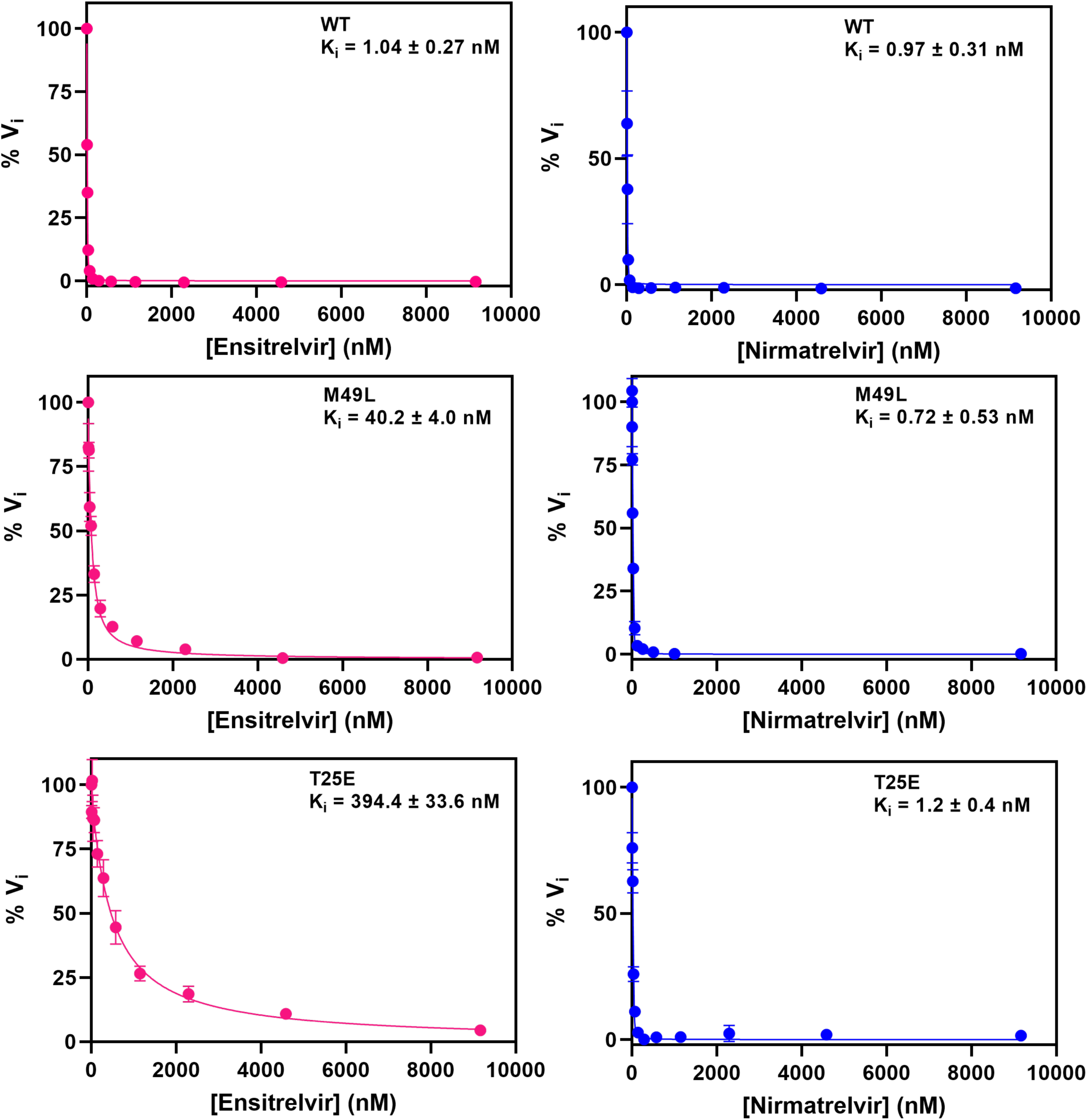
M49L and T25E Mpro exhibit decreased potency for ensitrelvir. Determination of Ki values for nirmatrelvir or ensitrelvir against Mpro using the FRET peptide assay. Curves were fit using the Morrison equation in Prism software. Each measurement was performed in triplicate.

There are four variants accessible by single nucleotide mutations that were in the top ten of DR scores for both nirmatrelvir and ensitrelvir (E166V, G2R, P168R, and Q299H), suggesting these variants may lead to cross resistance and are of particular health concern. The cross-resistant mutations indicate that given enough time nirmatrelvir and ensitrelvir are unlikely to provide a benefit in combination, necessitating the development of additional therapeutic tools to treat SARS-CoV-2 infections.

### Evaluation of resistance with regards to the substrate envelope

We explored the structural basis of variants likely to contribute to natural resistance by mapping them to the structure of M^pro^ and the substrate envelope. The substrate envelope represents the consensus volume occupied by the multiple M^pro^ cut sites in the viral proteome. Previous investigations of drug resistance demonstrate that inhibitors that fit within this consensus volume are less likely to be susceptible to drug resistance mutations. However, an inhibitor that protrudes beyond the substrate envelope provides opportunities for the evolution of resistance (Shaqra et al. 2022; Prabu-Jeyabalan, Nalivaika, and Schiffer 2002; Romano et al. 2011). Mutations that change the shape of the active site to fill sites of inhibitor protrusions can displace inhibitor while permitting substrate processing. We compared the substrate envelope with the location of bound inhibitors as well as the strongest apparent drug resistance mutations (Figure 6A). We focused on the strongest apparent drug resistance mutation here because steric clashes where inhibitors protrude from the substrate envelope typically have large impacts on inhibitor binding. Nirmatrelvir and ensitrelvir both protrude from the substrate envelope in the S1 pocket close to E166. Both inhibitors hydrogen bond to residue 166: nirmatrelvir to the side-chain and ensitrelvir to the main-chain.

**Figure 6.**
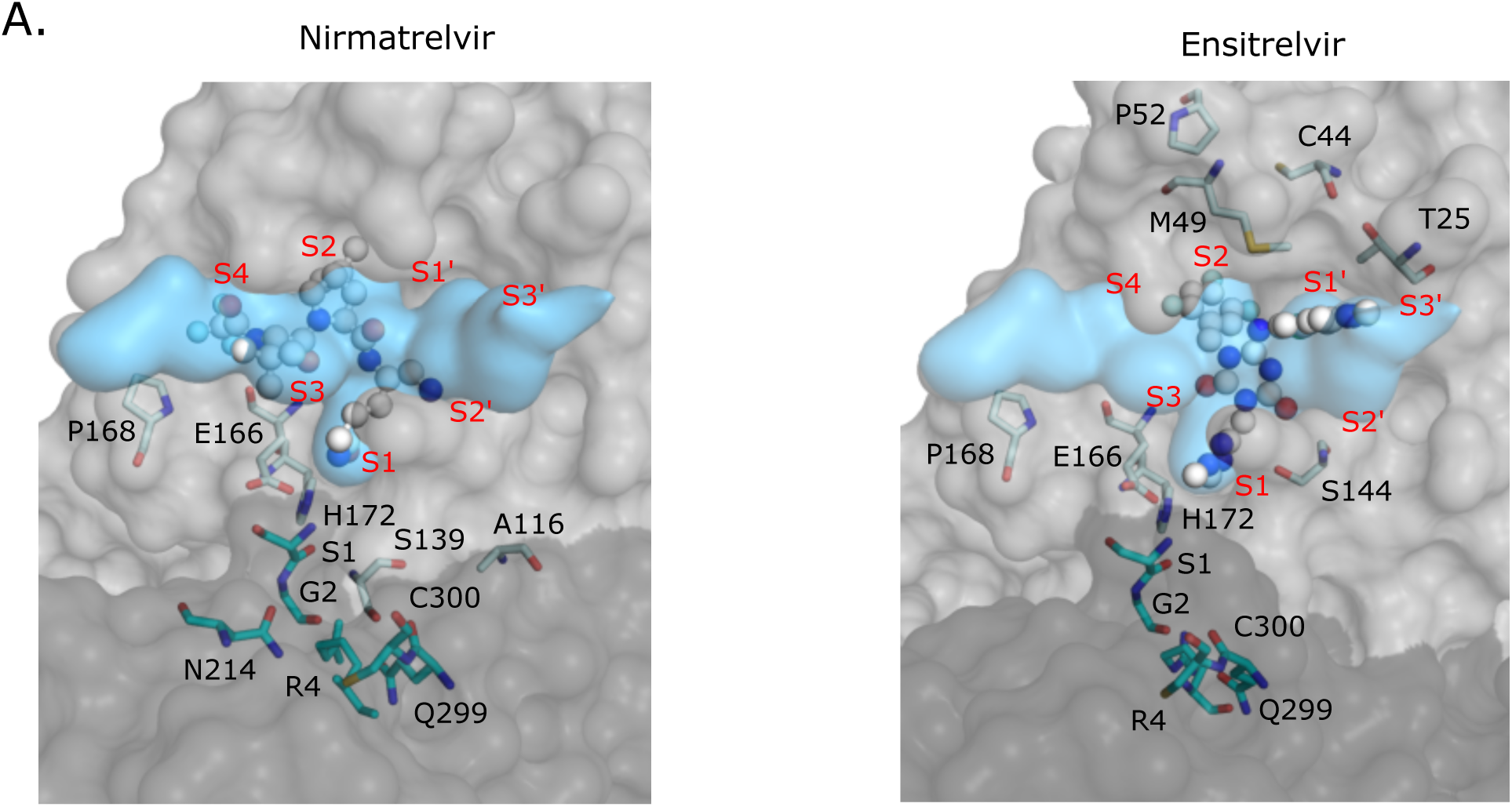
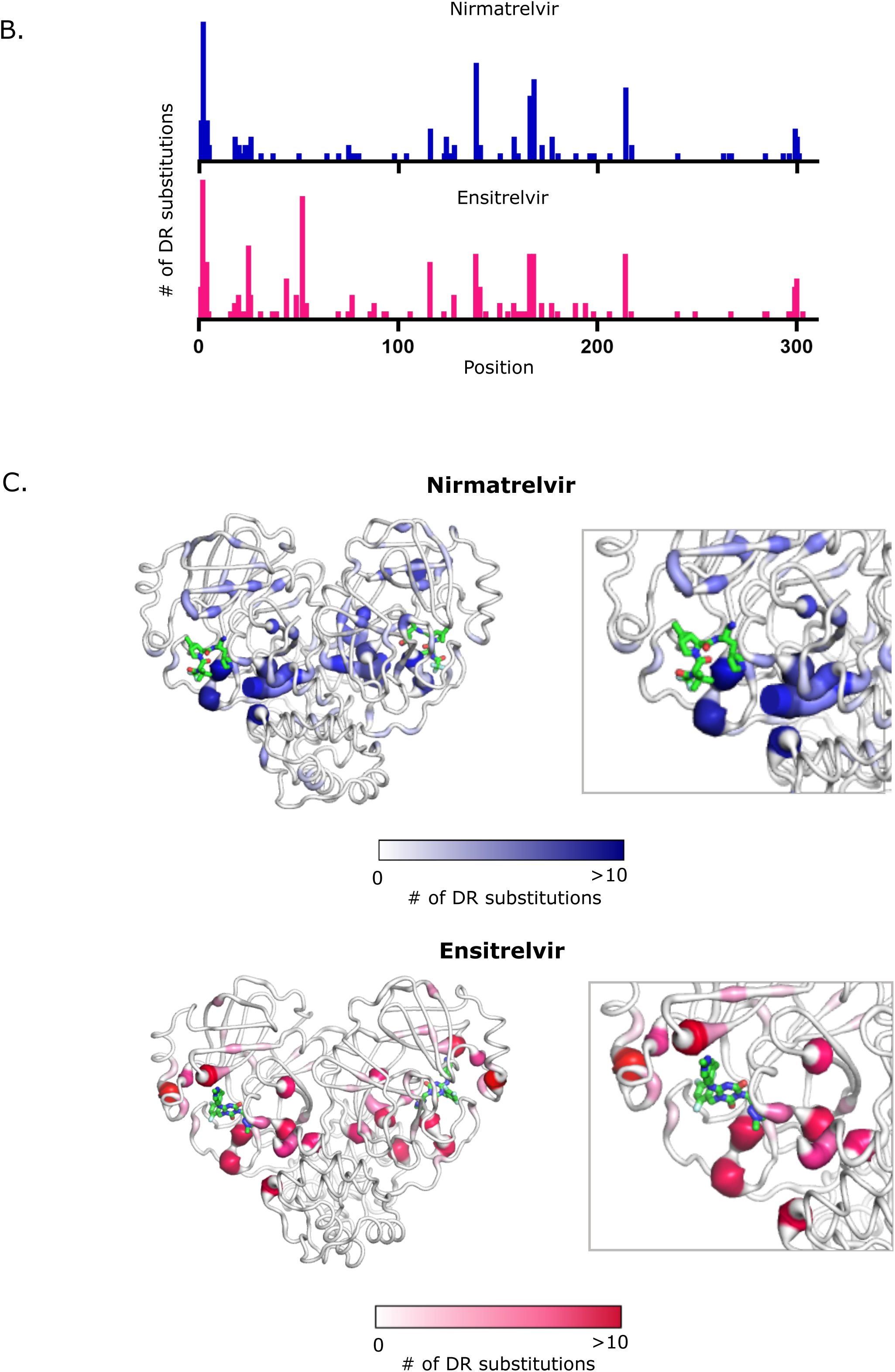
Structural distribution of apparent drug resistant mutations. A) The fit of nirmatrelvir (left panel) and ensitrelvir (right panel) within the substrate envelope. The labeled Mpro positions are those that contain the 30 mutations with the highest DR scores. S1, G2, R4, N214, Q299 and C300 are located on monomer B (shown in darker gray) and are represented as cyan sticks. The S4-S3’ binding subsites are labeled in red. B) The number of nirmatrelvir and ensitrelvir resistant mutations at each position of Mpro. C) The number of nirmatrelvir and ensitrelvir resistance mutations at each position of Mpro are mapped to the Mpro-inhibitor structures. Inhibitors are shown in green.

Mutations that cause the strongest cross-resistance to both inhibitors including G2R, E166V, P168R, and Q299H are located at or close to E166 where both nirmatrelvir and ensitrelvir protrude from the substrate envelope (Figure 6A). M^pro^ forms a dimer with two active sites. The active sites are formed predominantly by residues from one monomer, but the N-terminus of the trans-subunit extends close to the active site and forms a hydrogen bond with the side-chain of E166. G2 and Q299 are in steric contact with each other and mutations at either of these residues likely impact the dynamics and/or structure of the N-terminus of M^pro^ and the region of the active site near position 166. Similarly, P168 also appears positioned such that mutations are likely to impact the dynamics and/or structure of the active site in this region. P168 is in a beta-turn, where prolines are known to help stabilize or rigidify structure. Consistent with destabilization of this turn contributing to drug resistance, many other mutations at P168 also cause apparent drug resistance including K, V, and T.

### Inhibitor specific resistance mutations

We also observe some mutations that cause strong DR scores to one inhibitor but that did not impact the other inhibitor (Table 2). To explore the structural basis of inhibitor-specific resistance, we chose to investigate all amino acid changes regardless of the number of nucleotide mutations. The strongest inhibitor-specific resistance mutations were S1H and S139P for nirmatrelvir and mutations at C44, M49, T25, P52 and Y54 for ensitrelvir. Both position 1 and 139 are in proximity to E166 where both inhibitors protrude slightly from the substrate envelope. The amino terminus of position 1 hydrogen bonds to the side-chain of E166. S139 is in steric contact with positions 2 and 299 and is close to position 166. Many mutations at both position 1 and 139 (e.g, S1E, S1D, S139L and S139Q) show high DR scores for both inhibitors, consistent with similar protrusions from the substrate envelope closest to these residues. Thus, S1H and S139P may reshape the active site in a highly specific manner to preferentially disrupt interactions with nirmatrelvir. The strongest ensitrelvir-specific apparent resistance mutations including C44G, M49L, and P52L are all located close to unique protrusions of ensitrelvir from the substrate envelope in the S2 and S1’ pockets (Figure 6A). These results indicate that protrusions from the substrate envelope generally lead to resistance potential, though the drug resistance mutations themselves are challenging to predict from the substrate envelope alone.

**Table 2.**
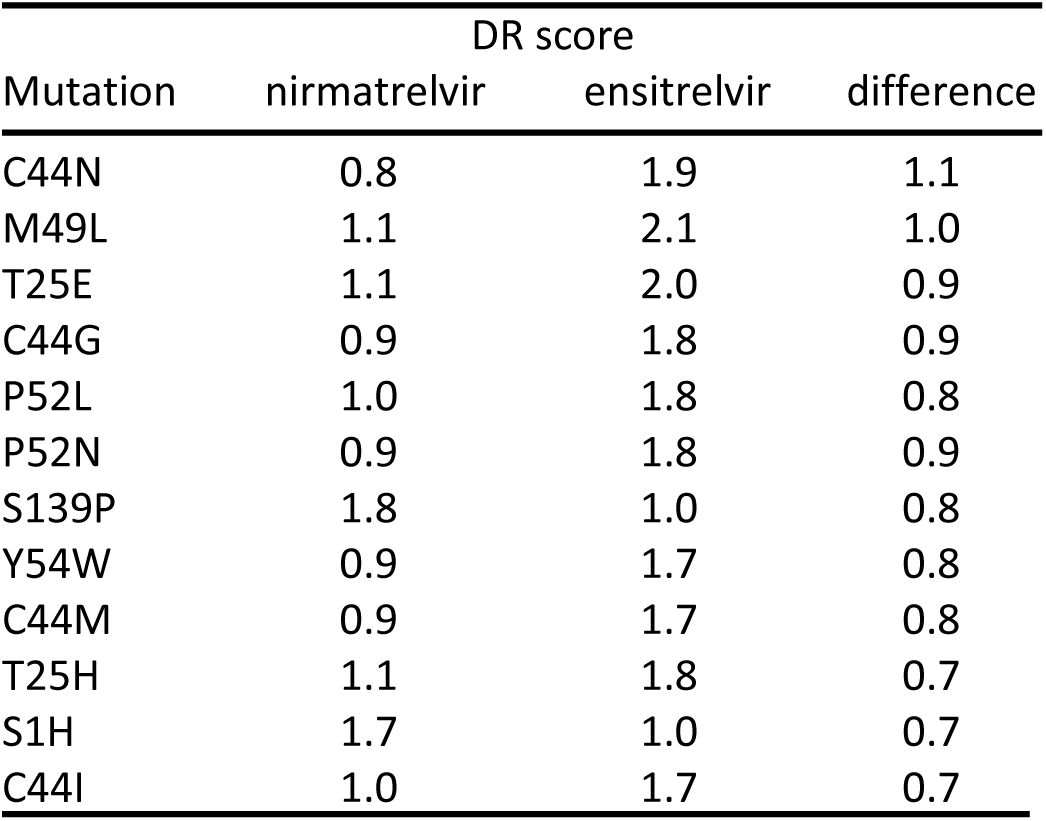
Mutations with largest differences in drug resistance score between nirmatrelvir and ensitrelvir.

| Mutation | DR score |  |  |
| --- | --- | --- | --- |
|  | nirmatrelvir | ensitrelvir | difference |
| C44N | 0.8 | 1.9 | 1.1 |
| M49L | 1.1 | 2.1 | 1.0 |
| T25E | 1.1 | 2.0 | 0.9 |
| C44G | 0.9 | 1.8 | 0.9 |
| P52L | 1.0 | 1.8 | 0.8 |
| P52N | 0.9 | 1.8 | 0.9 |
| S139P | 1.8 | 1.0 | 0.8 |
| Y54W | 0.9 | 1.7 | 0.8 |
| C44M | 0.9 | 1.7 | 0.8 |
| T25H | 1.1 | 1.8 | 0.7 |
| S1H | 1.7 | 1.0 | 0.7 |
| C44I | 1.0 | 1.7 | 0.7 |

### Location of all apparent drug resistance mutations provide mechanistic insights

To explore potential mechanisms of drug resistance, we mapped all apparent resistance mutations to structure (Figure 6B and C). Apparent drug resistance mutations were identified at 63 different positions in M^pro^ (Supplementary Table 2). However, the majority (190 of 319) of the apparent drug resistance mutations occurred at 14 hot-spot positions (Table 3). The mutations driving the largest effects (e.g., Table 1) are almost all at these hot-spot positions. The main exceptions are H172Y (DR score of 1.9 for ensitrelvir) and M49L (DR score of 2.1 for ensitrelvir). Both of these positions were on the fringe of the cutoff we used for hot-spots (6 apparent DR mutations). At position 172 the only tolerated mutations that result in enzymatic function in our assay are aromatic amino acids, and indeed H172Y and H172F were apparent DR mutations for both inhibitors. At M49, three mutations (L, I, H) caused apparent DR for ensitrelvir. Position 49 is located close to unique protrusions of ensitrelvir from the substrate envelope and there were no apparent DR mutations for nirmatrelvir at this position. Upon examination of the structure of M^pro^, all of the hot-spot positions were located at or adjacent (via bridging contacts with one reside) to the active site and/or the dimerization interface (Table 3).

**Table 3.** Positions with the most apparent drug resistance mutations.

| Position | Apparent DR mutations |  | Location |
| --- | --- | --- | --- |
|  | Nirmatrelvir | Ensitrelvir |  |
| G2 | 17 | 17 | Dimer interface adjacent to active site |
| S139 | 12 | 8 | Dimer interface adjacent to active site |
| P168 | 10 | 8 | Active site and dimer interface |
| N214 | 9 | 8 | Adjacent to dimer interface |
| E166 | 8 | 8 | Active site and dimer interface |
| P52 | 0 | 15 | Adjacent to active site |
| R4 | 5 | 7 | Dimer interface |
| A116 | 5 | 7 | Adjacent to active site and dimer interface |
| S1 | 5 | 4 | Dimer interface adjacent to active site |
| T25 | 0 | 9 | Active site |
| Q299 | 4 | 4 | Dimer interface |
| C300 | 3 | 5 | Dimer interface |
| T26 | 3 | 3 | Active site |
| L141 | 2 | 4 | Dimer interface adjacent to active site |

These observations indicate that the dynamics and/or structure of the active site is coupled to the structure of the dimer interface. The majority of the indirect resistance hot-spots for both inhibitors including positions 1, 2, 4, 214, 298 and 299 contact one another and span from one active site across the dimer interface to the second active-site (Figure 6C). In our previous functional scan of M^pro^ without inhibitors (Flynn et al. 2022) we noted a hypersensitive amino acid communication network spanning between active sites that we proposed modulated the shape and/or dynamics of the active sites. The resistance hot-spot positions provide further evidence that dimerization can strongly influence the physical properties of the active site. Of note, dimerization of M^pro^ is essential for its catalytic activity (Tan et al. 2005; Shi, Sivaraman, and Song 2008). In addition, mutations at the substrate binding site including E166A (Cheng, Chang, and Chou 2010) and S147A (Barrila, Bacha, and Freire 2006) weaken M^pro^ dimerization, consistent with the shape of the active site being linked to dimerization. Structurally, the two M^pro^ subunits are oriented perpendicular to each other with an extensive interface formed primarily between domain II of one monomer and the N-terminal residues (N-finger) of the other monomer (Zhang et al. 2020). Direct interaction between the N-finger from one subunit and E166 at the active site in the other subunit have been shown to strongly impact M^pro^ activity (Cheng, Chang, and Chou 2010). Our comprehensive drug resistance study demonstrates that changes in a network of amino acids that bridge between the active sites can disrupt binding to both nirmatrelvir and ensitrelvir.

### Functional cost of apparent drug resistance mutations and implications for evolution

Both M^pro^ function and affinity for inhibitor may contribute to the evolution of drug resistance in circulating viral populations. M^pro^ variants with wild-type-like function that strongly disrupt inhibitor binding pose the greatest chance of increasing to high frequency and causing the most harm to human health. We examined the relationship between DR score and functional score (Figure 7). As DR scores increase, there is a clear trend towards lower functional scores. Mutations that cause the strongest disruption of drug binding tend to also disrupt the processing of substrates. Indeed, only one amino acid change accessible by a single nucleotide change (P168R for nirmatrelvir) led to a DR score greater than 1.8 and a functional score equal to or greater than wild-type. The rarity of this type of mutation increases the chance of evolving resistance through alternative pathways that require multiple mutations as has been observed in SARS-CoV-2 passaging studies (Iketani et al. 2022a). This is similar to what has been observed for the evolution of drug resistance in HIV to protease inhibitors. Primary resistance mutations that disrupt inhibitor binding to HIV protease typically possess reduced enzyme activity. Continued drug pressure results in the emergence of compensatory secondary mutations that rescue these enzymatic defects (Liu et al. 2005; Kozisek et al. 2007; Schock, Garsky, and Kuo 1996).

**Figure 7.**
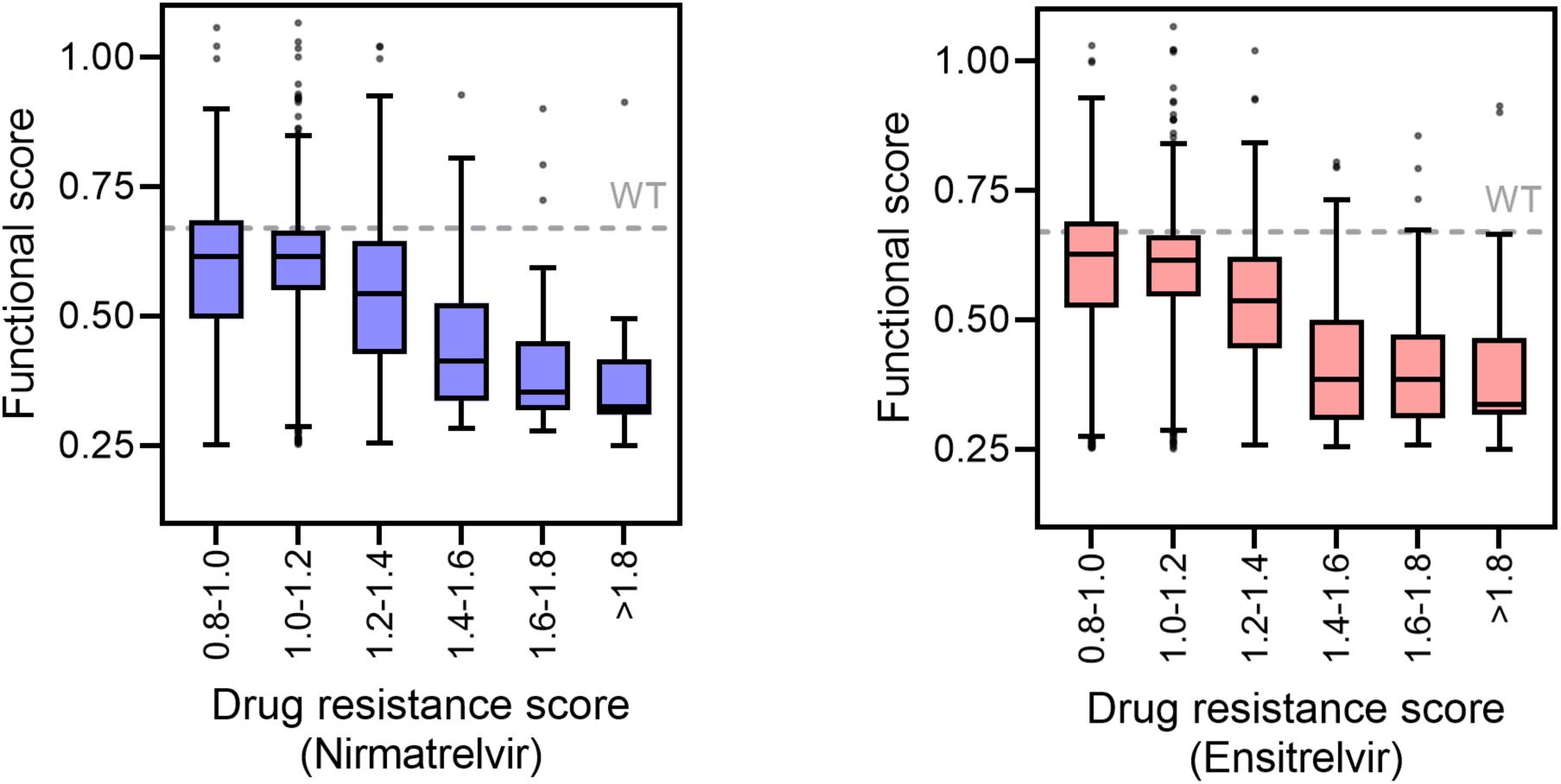
Functional cost of apparent drug resistant mutations. Mutants with increased DR scores for nirmatrelvir (left panel) or ensitrelvir (right panel) tend to have reduced functional scores. The box and whisker plots are plotted using the Tukey method. The box contains the 25th to 75th percentiles of the dataset with the center line denoting the median value.

In our data, we identified a large group of mutations that increase the enzyme activity of M^pro^ (Supplemental table 1). Due to the strong reproducibility of our functional scores, some mutations with small increases in function were statistically significant in these analyses. The large decreases in function of variants with the strongest DR scores (Figure 7) should select for compensatory mutations with large impacts on function. For example, T21I and L50F were observed multiple times in viral selection experiments with nirmatrelvir (Iketani et al. 2022a; Jochmans et al. 2023; Zhou et al. 2022) and both of these variants exhibit strong increases in functional score (0.92 and 0.89 for T21I and L50F respectively compared to 0.67 for wild-type). Because enzyme function can depend on substrate, we are pursuing analyses of increased activity mutants using substrates directly relevant to viral infections. A thorough analysis of activating mutations is beyond the scope of this work and will be the topic of future efforts.

### Survey of drug resistance in natural isolates

Our results as well as those from passaging studies indicate that clinical use of drugs targeting M^pro^ will select for multiple different mutational pathways. Therefore, surveillance efforts for detecting early steps in drug resistance evolution will be more powerful if they integrate over all likely pathways to resistance. Our comprehensive survey of mutations that can cause apparent drug resistance provides a promising approach to this issue. We surveyed for all variants that we identify as likely to contribute to drug resistance based on DR score and accessibility by single nucleotide mutations. We also assessed the frequency of these mutations in SARS-CoV-2 isolates obtained in the U.S. both before and after approval of Paxlovid (data not shown). These initial analyses indicated that resistance in circulating virus populations is currently low and/or undetectable. The reproducible measurements of drug resistance potential described here provide a valuable tool for future efforts to assess resistance in data from ongoing sequencing efforts. With this motivation, we are currently working towards developing publicly available computational tools to survey the frequency of variants with either strong DR scores and/or high functional scores as a function of geographic location and time relative to clinical use of M^pro^ inhibitors.

### Caveats and sources of uncertainty

Simplified readouts of the impacts of mutations such as those presented in this work are not perfect mimics of impacts in viruses. For example, we analyzed M^pro^ mutations expressed in yeast, which provides a distinct environment including distinct chaperones compared to human hosts. While some proteins are highly sensitive to chaperones for folding, M^pro^ can be readily expressed and folded in bacteria (Zhang et al. 2020). In addition, M^pro^ is active in purified form indicating that it is not highly sensitive to chaperones or other cellular factors for function. We used yeast growth as a readout of M^pro^ function that we assume is caused by cleavage of endogenous yeast proteins that are not directly related to viral infection. Because our analyses of drug resistance score are determined by changes in function with and without drug, the impact of using non-viral substrates is mitigated as inhibitors should impact all substrates similarly. Indeed, independent measures of IC_50_’s of M^pro^ variants in purified form correlate with our drug resistance scores (Supplemental Figure 8) (Hu et al. 2022). For assessment of mutant effects on M^pro^ function (e.g., increased activity mutations), it may be more important to use cut-sites from SARS-CoV-2, and we are actively investigating how mutations can increase M^pro^ function using cleavage sites that are native to SARS-CoV-2.

While it is clear that M^pro^ function is required for SARS-CoV-2 infectivity, the quantitative relationship between function and fitness as not been accurately assessed. Thus, we do not know how viral fitness is determined based on the activity of M^pro^ nor inhibitor concentration and affinity. Because there are numerous viral cut-sites, developing an accurate model of relationships between fitness and M^pro^ function is complex. In the absence of an accurate quantitative model, we can still make qualitative assessments. For example, there is strong evidence that M^pro^ mutants with very low functional scores are selected against in viral populations (Flynn et al. 2022) and that mutations that reduce binding affinity for inhibitors increase the ability of virus to expand in the presence of drugs (Iketani et al. 2022b). Thus, the evolution of M^pro^ variants with wild-type-like function and decreased affinity for inhibitors clearly represents an undesirable outcome – a virus that would be both highly drug resistant and highly fit in the absence of drug. For viral maturation, M^pro^ must cut itself out of the viral polyprotein such that sequences at the N-terminus and C-terminus of M^pro^ may have special relevance in the context of viral replication that are not captured in our screen. Therefore, the viral relevance of DR scores for M^pro^ mutations near the termini should be interpreted with caution.

Our approach describes point mutations only and therefore does not provide information on the potential interdependence of mutations at multiple positions in the M^pro^ protein. Our observations and those from others (Iketani et al. 2022b; Jochmans et al. 2023; Zhou et al. 2022) indicate that the natural evolution of high levels of drug resistance is likely to involve multiple mutations including primary mutations that strongly disrupt drug binding (but come with functional costs) and compensatory mutations that rescue function. Our data provides a reliable view of mutations that disrupt drug binding and a first pass at mutations that increase function in the wild-type M^pro^ genetic background. In considering multiple mutations, independent or near-independent effects of mutations can be a reasonable first approximation (Sternke, Tripp, and Barrick 2019), but epistasis or mutational dependencies are also a common feature of protein evolution (Pollock and Goldstein 2014). In future work, it will be important to evaluate the contributions of epistasis to M^pro^ evolution.

## Conclusions

Many mutations in M^pro^ can cause cross-resistance to both nirmatrelvir and ensitrelvir, providing strong motivation for the development of additional therapeutics to combat SARS-CoV-2 infections. Mutations that can cause cross-resistance were predominantly located at or near E166. Both nirmatrelvir and ensitrelvir protrude slightly from the substrate envelope to make hydrogen bonds to either the side-chain or main-chain of E166. The close contact of nirmatrelvir and ensitrelvir with E166 appear to make them susceptible to similar resistance mutations. From the perspective of resistance potential, these findings provide motivation for developing inhibitors that do not hydrogen bond with E166 and bind within the substrate envelope and/or inhibitors that target other sites or other proteins as has been successful in treatments for HIV (Shafer and Vuitton 1999).

Mutations with the largest DR scores tend to reduce the enzymatic activity of M^pro^ such that drug pressure is likely to select for a combination of primary resistance mutations that reduce function together with compensatory mutations that rescue activity. Because many mutations cause either drug resistance or increased function, there are a wide array of variants likely to evolve in response to drug pressure. The comprehensive identification of resistance mutations of interest provides for sensitive and thorough surveillance in the early stages of evolution of drug resistance in M^pro^. Variants with strong drug resistance together with wild-type-like enzymatic function provide the greatest potential to expand to high prevalence with significant impacts to public health. This pattern of resistance evolution is reminiscent of recent observations in influenza A virus (Abed et al. 2011), where selective pressure against oseltamivir caused the evolution of a primary neuraminidase mutation that disrupted inhibitor binding together with secondary mutations that rescued function. The resulting fit and resistant neuraminidase variant rose in frequency until it was dominant in viral isolates in 2008 (Hurt et al. 2011). The frequency of resistant variants of M^pro^ in SARS-CoV-2 isolates may be used as a barometer for the likelihood of widespread resistance evolution.

## Materials and Methods

### Generation of mutant libraries and barcode association

The SARS-CoV-2 single site variant library was synthesized by Twist Biosciences (twistbioscience.com) with each amino acid position modified to all 19 amino acid positions plus a stop codon using the preferred yeast codon for each substitution. The library was fused to a Ubiquitin gene, combined via Gibson Assembly with a linearized p416LexA destination vector, and barcoded with a randomized 18 bp nucleotide sequence as described in Flynn et al, 2022 (Flynn et al. 2022). The p416 LexA destination vector contains a LexA-ER-B112 transcription factor which is activated by β-estradiol (Ottoz, Rudolf, and Stelling 2014).

### Bulk competition experiment of M^pro^ libraries in presence of inhibitors

The barcoded plasmid library was combined with a plasmid containing wild-type M^pro^ linked to approximately 150 unique barcodes (Flynn et al. 2022) and was transformed using the large-scale lithium acetate procedure (Gietz and Schiestl 2007) into CMB855 cells (*Mat a ura3 leu2 his3 lys2 snq2::kanMX pdr5::kanMX pdr1::nat1 yap1::nat1*). Sufficient transformation reactions were performed to attain 2 million independent yeast transformants representing a 20-fold sampling of the average barcode. Following a 6-hour recovery in YPAD, transformed cells were washed three times in SD-U media (synthetic dextrose media lacking uracil to select for the plasmid library) to remove extracellular DNA and grown in 1 L SD-U at 30°C for 40 hours with repeated dilution to maintain the cells in log phase of growth (OD_600_ = 0.05 - 1) and to expand the library. The library was diluted to early log phase in SD-U, grown for 3 hours, and a 0-hour time point sample of 10^8^ cells was collected by centrifugation and cell pellets were stored at - 80°C. Cells were subsequently split into flasks containing 10 mL each, and either 10 µM nirmatrelvir (from a 50 mM stock in 100% DMSO), 20 uM ensitrelvir (from a 50 mM stock in 100% DMSO) or 2 µL of 100% DMSO was added to each flask. Additionally, 0.005% SDS was added to permeabilize the cell membrane in order to increase uptake of the drug. To analyze the accuracy of the growth experiments, we performed three technical replicates with each inhibitor and the DMSO control. For replicates, we transformed the library into yeast cells in bulk, performed an outgrowth, then separated the culture into three flasks and performed the growth competitions and sequence analyses independently. Each drug condition was performed in triplicate. 2 µM β-estradiol (from a 10 mM stock in 95% ethanol) was then added to each flask to activate expression of M^pro^. Cultures were grown with shaking at 180 rpm for 16 hours with dilution after 8 hours (into media containing the appropriate concentrations of β-estradiol, nirmatrelvir, ensitrelvir, DMSO, and SDS). 16-hour samples were collected by centrifugation and cell pellets were stored at -80°C.

### Determination of functional scores

DNA was isolated, amplified and the barcodes were sequenced on an Illumina NextSeq instrument described (Flynn et al. 2022) with minor modifications. Purified plasmid DNA from each time point was linearized with AscI. Barcodes were amplified with 22 cycles of PCR using Phusion polymerase (NEB) with primers that add Illumina adaptor sequences and a 6 bp identifier sequence to distinguish the various drug conditions and time points. PCR products were purified two times over silica columns (Zymo research) and DNA concentration was quantified using the KAPA SYBR FAST qPCR Master Mix (Kapa Biosystems) on a BioRad CFX Machine. PCR samples were combined and sequenced on an Illumina NextSeq instrument using a NextSeq 500/550 High Output Kit v.2.5 (75 cycles). The Illumina barcode reads were calculated using custom scripts that have been deposited on Github (https://github.com/JuliaFlynn/BolonLab) as previously described (Flynn et al. 2022). Illumina sequences were filtered for reads with a Phred score >10 and strict matching to the expected template and identifier sequence. Filtered reads were parsed based on their identifier sequences. For each condition/time point each unique N18 barcode read was counted. Using the variant-barcode association table that was generated by PacBio sequencing (see (Flynn et al. 2022)) the unique N18 count file was used to calculate the frequency of each mutant in each condition/time point. The abundance of each mutant at each time point was tabulated and normalized to the total number of stop reads for amino acids 1-299. Stop codon reads for amino acids 300-306 were excluded because M^pro^ mutants containing stops at these positions remain functional (Flynn et al. 2022). Functional scores were determined by linear fits to the change in mutant abundance relative to stop mutations. All figures in this paper use the average functional score between the three replicates. The standard deviation was used to represent experimental variation between replicates. Drug resistance scores were calculated as the ratio of the average functional score of each mutant in the absence of the inhibitor to that in the presence of the inhibitor, compared to the same ratio of wild-type M^pro^. The standard deviations between the functional scores in the three replicates with and without drug were propagated to assess statistical significance and calculate p-values for the drug resistance scores.

### Purification of M^pro^

His_6_-SUMO-SARS-CoV-2 M^pro^ (WT) was cloned into a pETite vector (Shaqra et al. 2022). M49L and T25E mutations were created by site specific mutagenesis. The vectors were transformed into Hi-Control BL21(DE3) *E. coli* cells and the M^pro^ proteins were expressed and purified as previously described (Shaqra et al. 2022) with minor modifications. An overnight culture was used to inoculate 1 L of Terrific Broth (TB) medium containing 50 µg/ml of kanamycin at 37°C. Once the culture reached OD_600_ = 1.5, protein expression was initiated with addition of 0.5 mM IPTG, the temperature was reduced to 19°C, and cells were grown overnight. Cells were harvested by centrifugation and resuspended in lysis buffer containing 50 mM Tris pH 8.0, 400 mM NaCl, 1mM TCEP prior to lysis by two passes through a cell disrupter at 15,400 psi. The lysed cell suspension was clarified by centrifugation at 19,000 rpm for 30 min at 4°C and the supernatant was incubated with Co-NTA resin that had been pre-equilibrated with lysis buffer for 2 hours at room temperature. The following steps were all performed at room temperature due to a proclivity for the protein to precipitate at 4°C. The resin was loaded onto a column and was washed with lysis buffer containing 10 mM imidazole until no protein was detected in the flow through (about 20 column volumes). M^pro^ was eluted with 5 ml of lysis buffer containing 500 mM Imidazole. The SUMO tag was then cleaved by addition of Ulp1 to the eluted protein while dialyzing into 2 L of lysis buffer in a 10,000 MWCO dialysis cassette overnight. Following digestion, the SUMO protease and SUMO tag were removed by CoNTA resin. Protein was concentrated to 2.5 ml and run over a Sephadex G-25 desalting column (Cytiva) pre-equilibrated with 25 mM HEPES pH 7.5, 150 mM NaCl, 1 mM TCEP. Proteins were frozen in liquid nitrogen and stored at -80°C.

### Determination of Michaels-Menten (K_m_) constant

To measure the K_m_ of M^pro^ mutants, 100 nM enzyme was added to a series of 0-125 µM FRET substrate (Dabcyl-KTSAVLQSGFRKME-Edans (LifeTein)) in assay buffer (50 mM Tris pH 7.5, 50 mM NaCl, 1 mM EDTA, 1 mM DTT and 4% DMSO). The proteolytic reaction was monitored using a Perkin Elmer EnVision plate reader (excitation at 355 nm, emission at 460 nm). Three replicates were performed with each mutant. The initial velocity was plotted against FRET substrate concentrations and were fit using GraphPad Prism 9 to the Michaelis-Menten equation.

### Enzyme inhibition assays

For K_i_ measurements, mutant M^pro^ enzyme was pre-incubated at 30°C with increasing concentrations of ensitrelvir or nirmatrelvir for 45 min in assay buffer. The proteolytic reaction was initiated with a final concentration of 10 µM protease FRET substrate (LifeTein) and monitored using a Perkin Elmer Envision plate reader (excitation at 355 nm, emission at 460 nm). Three replicates were performed for each mutant and each inhibitor. The reaction was monitored for 1 hour and the initial velocity was calculated by linear regression. The K_i_ was calculated by plotting the inhibitor concentration versus the initial velocity and fit to the Morrison equation (tight binding) using GraphPad Prism 9 software.

### van der Waals calculations

All crystal structures were obtained from the Protein DataBank. Prior to van der Waals calculations, M^pro^-inhibitor complexes were prepared using the Schrodinger Protein Preparation Wizard. Hydrogen atoms were added, protonation states determined using PropKa, and the hydrogen bonding network was optimized. A restrained minimization was performed using the OPLS2005 force field within an RMSD of 0.3 Å. All crystallographic waters were retained during structure minimization. Interaction energies between the inhibitor and protease were estimated using a simplified Lennard-Jones potential using a custom Python script, as previously described.

### PCA analysis

Prior to analysis, all vdW interactions were standardized by calculating the Z score (Equation 1); the sample mean was subtracted from each interaction and divided by the standard deviation. The Z scores were used as input for the scikit-learn implementation of principal components analysis. Principal components accounting for 65% of the variance were used to reproject the vdW data, and we used k-means clustering to separate inhibitors into three clusters.

### Structure analysis

The inhibitor co-crystal structure (PDB 7vu6 for ensitrelvir (Tyndall 2022) and PDB 7RFS for nirmatrelvir (Owen et al. 2021) were aligned onto the co-crystal structure of SARS-CoV-2 M^pro^ with the Nsp4/5 substrate peptide (PDB 7T70) using the carbon alpha atoms of residues 41, 144, 145, 163 and 164 (Shaqra et al. 2022). This allows for proper visualization of each inhibitor within the substrate envelope. Because 7vu6 lacks the two N-terminal residues S1 and G2, these were modeled in using the conformations seen in PDB 6xhm as a reference (Hoffman et al. 2020). Figures were generated using Matplotlib (Hunter 2007), PyMOL, and GraphPad Prism version 9.3.1.

### Data availability

Next generation sequencing data has been deposited to the NCBI short read archive (PRJNA933538).

---

**Supplemental Figure 3.**
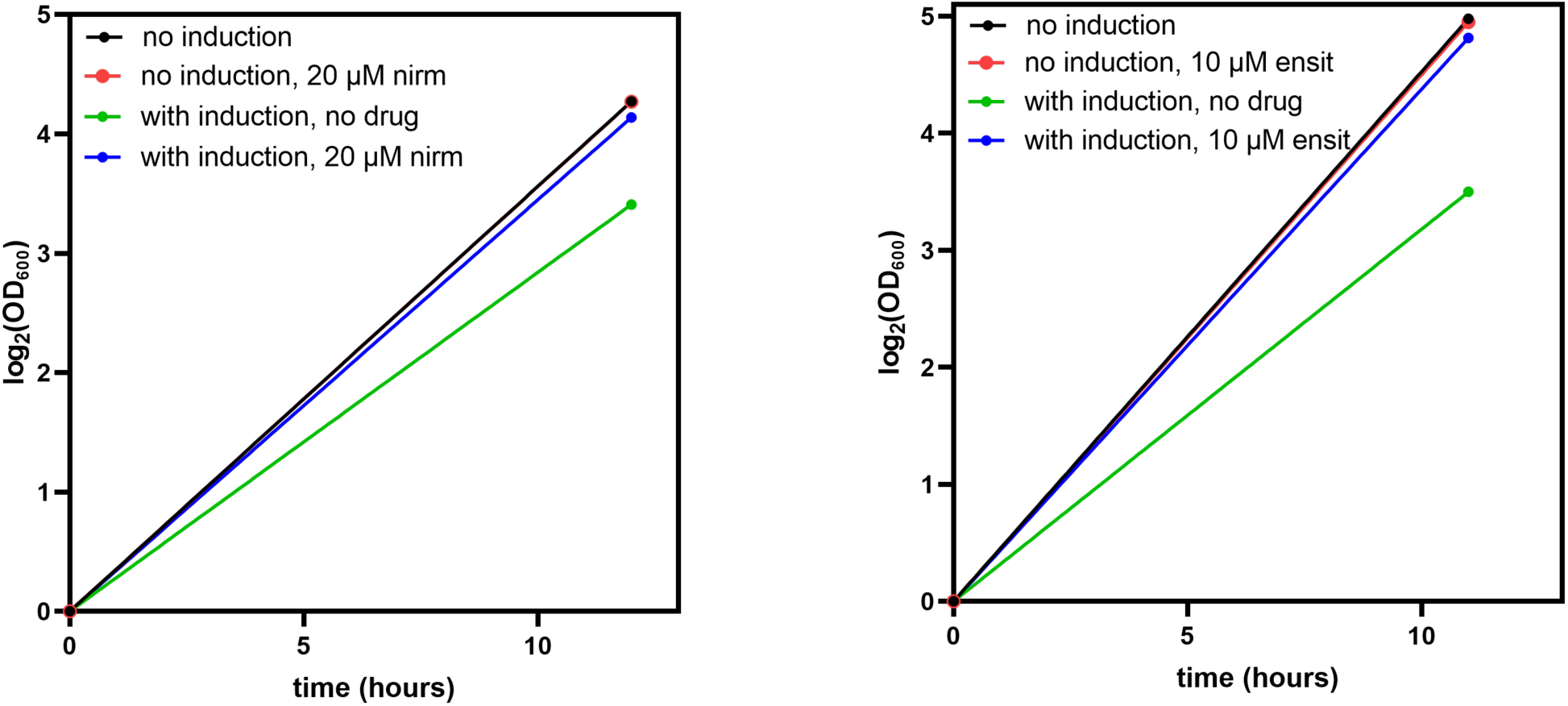
Toxicity of Mpro to yeast cells is inhibited by nirmatrelvir and ensitrelvir. The growth rate of Δ4 cells harboring a plasmid encoding for wild-type Mpro was measured in the absence or presence of 2 µM β-estradiol to induce Mpro expression and either nirmatrelvir (left panel) or ensitrelvir (right panel).

**Supplemental Figure 4.**
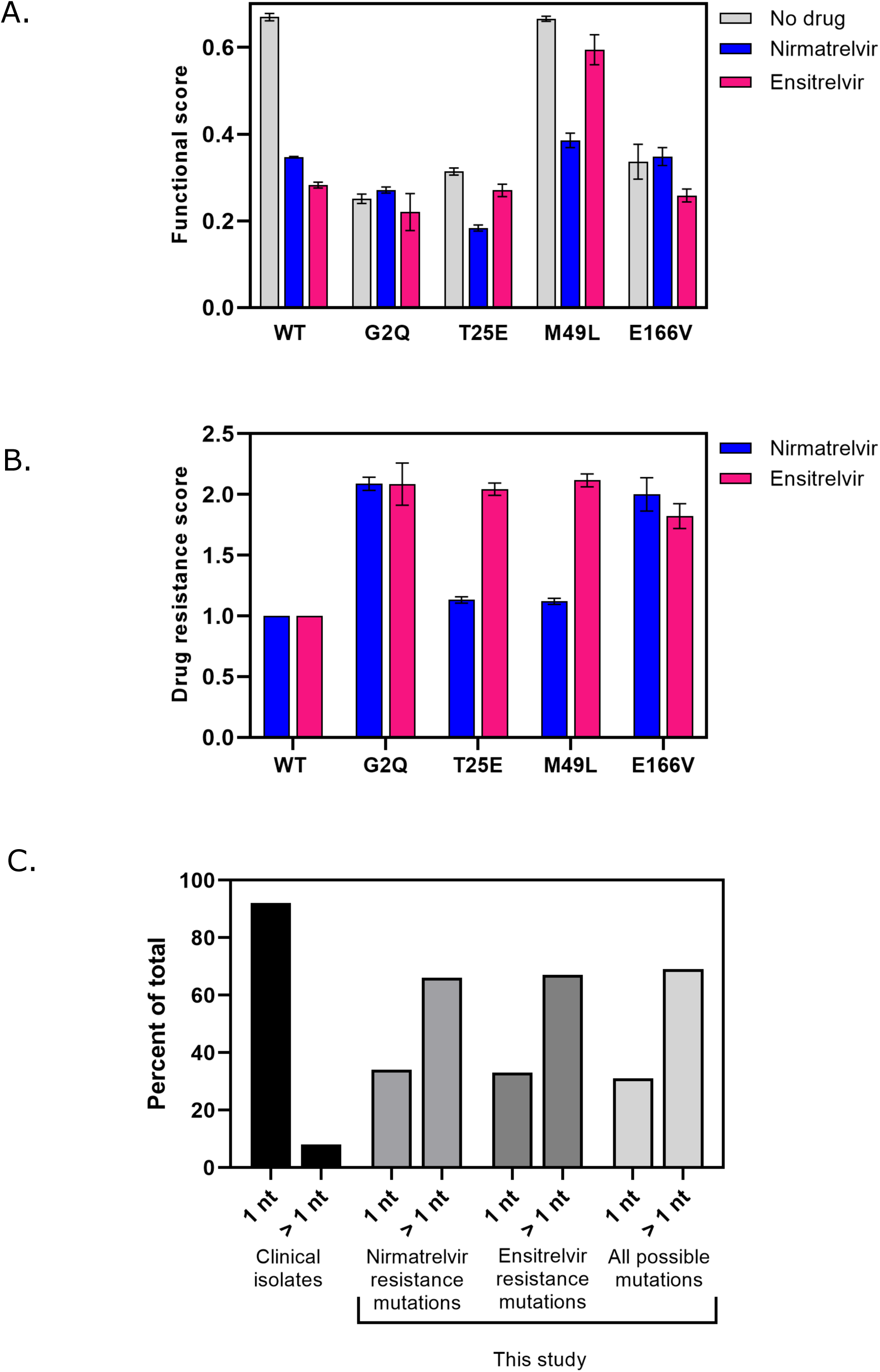
A) The functional score of certain library members are less sensitive to inhibitors. B) Calculated DR scores for various Mpro mutants against nirmatrelvir and ensitrelvir. C) The number of nucleotide mutations to make drug resistance mutations in this study as compared to clinically identified variants.

**Supplemental Figure 5.**
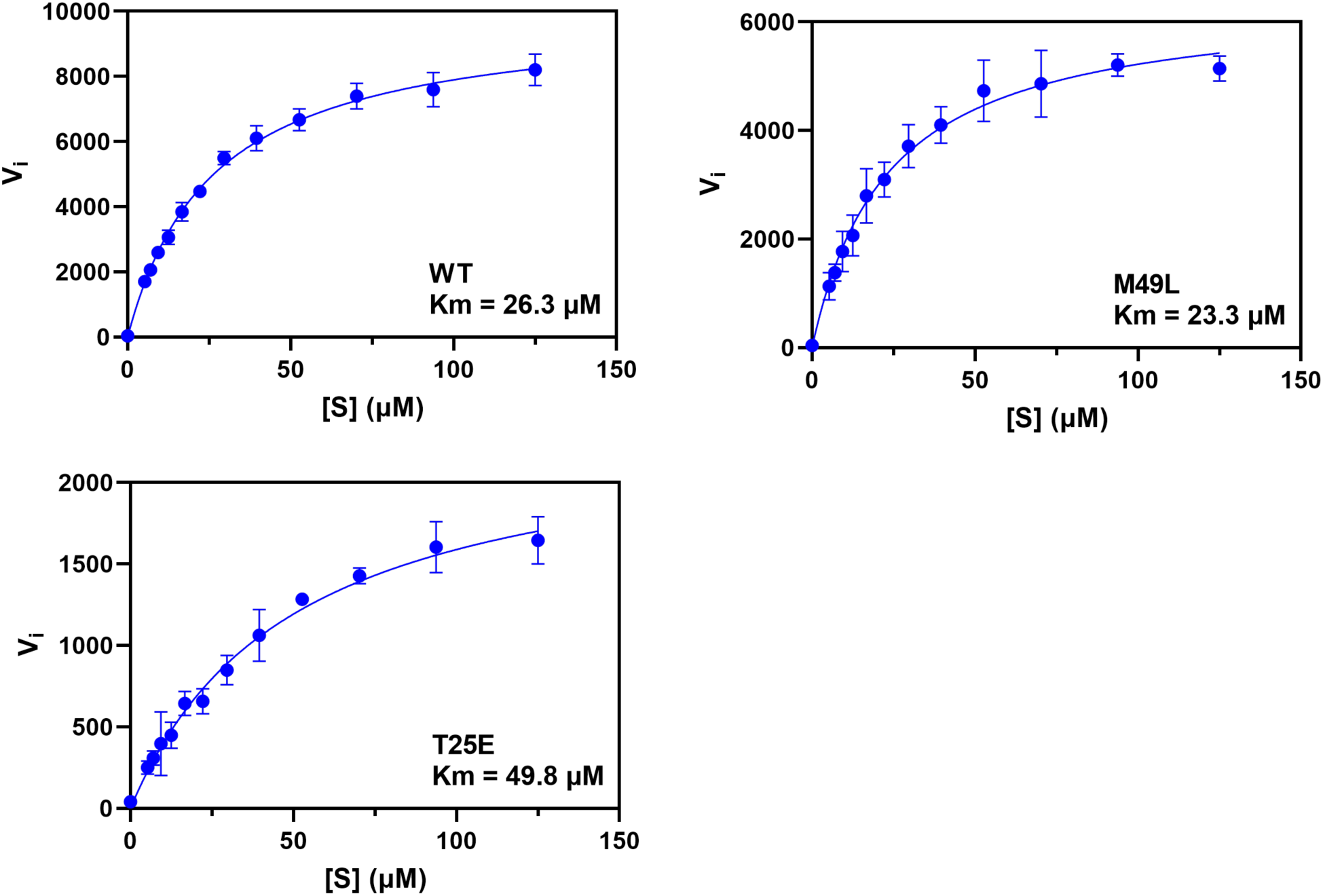
Enzymatic characterization of purified Mpro mutants.

**Supplemental Figure 8.**
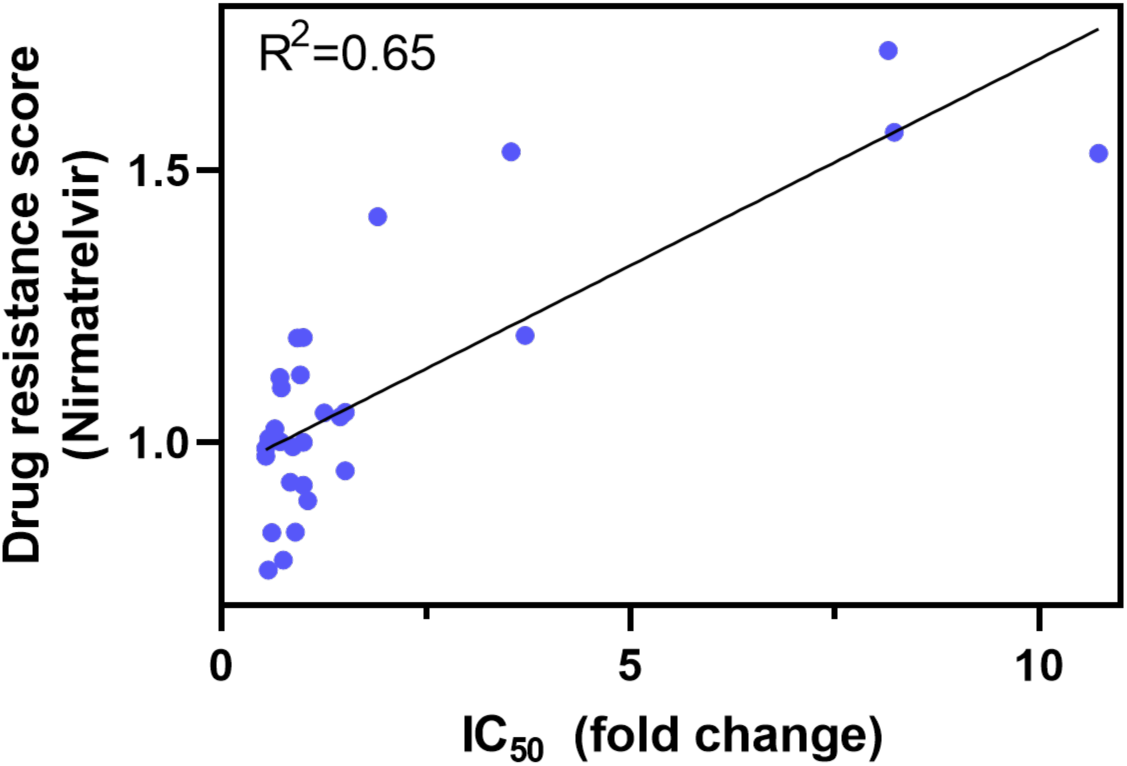
IC50s of purified Mpro variants in purified form correlate with our drug resistance scores. IC50 values of select Mpro mutants were measured by Hu et al (Hu, Lewandowski et al. 2022).

